# Asymmetric triangular body-cover model of the VFs with bilateral intrinsic muscle activation

**DOI:** 10.1101/2024.03.18.585590

**Authors:** Jesús Parra, Carlos Calvache, Gabriel Alzamendi, Emiro Ibarra, Leonardo Soláque, Sean D. Peterson, Matías Zañartu

**Affiliations:** Department of Electronic Engineering, Universidad Técnica Federico Santa Maria, Valparaíso, Chile; Department of Mechatronics Engineering, Universidad Militar, Bogotá, Colombia; Department Communication Sciences and Disorders, Corporación Universitaria Iberoamericana, Bogotá, Colombia; Vocology Center, Bogotá, Colombia; Institute for Research and Development on Bioengineering and Bioinformatics (IBB), CONICET-UNER, Oro Verde, Entre Ríos 3100, Argentina; Facultad de Ingeniería, Universidad Nacional de Entre Ríos, Entre Ríos, Argentina; Department of Mechanical and Mechatronics Engineering, University of Waterloo, Waterloo, Ontario, Canada

## Abstract

Many voice disorders are linked to imbalanced muscle activity and known to exhibit asymmetric vocal fold vibration. However, the relation between imbalanced muscle activation and asymmetric vocal fold vibration is not well understood. This study introduces an asymmetric triangular body-cover model of the vocal folds, controlled by the activation of intrinsic laryngeal muscles, to investigate the effects of muscle imbalance on vocal fold oscillation. Various scenarios were considered, encompassing imbalance in individual muscles and muscle pairs, as well as accounting for asymmetry in lumped element parameters. The results highlight the antagonistic effect between the thyroarytenoid and cricothyroid muscles on the elastic and mass components of the vocal folds, as well as the impact on the vocal process from the imbalance in the lateral cricoarytenoid and interarytenoid adductor muscles. Measurements of amplitude and phase asymmetry were employed to emulate the oscillatory behavior of two pathological cases: unilateral paralysis and muscle tension dysphonia. The resulting simulations exhibit muscle imbalance consistent with expectations in the composition of these voice disorders, yielding asymmetries exceeding 30% for paralysis and below 5% for dysphonia. This underscores the versatility of muscle imbalance in representing phonatory scenarios and its potential for characterizing asymmetry in vocal fold vibration.

## I. INTRODUCTION

The production of human communication sounds results from intricate physical interactions within the speech organs, governed by a sophisticated neuromuscular control system (Story, 2015; Titze and Story, 2002). Vocal physiology can be elucidated through (1) the nearly stable subglottal driving pressure, (2) the intricate interplay of sound, flow, and vocal fold (VF) tissue at the glottal level, and (3) the propagation and modulation of voice sound throughout the supraglottal vocal tract (Story, 2002; Titze and Hunter, 2007; Zhang, 2016). A diverse array of numerical models exists, varying in physiological fidelity, mathematical intricacy, and computational demands (Story, 2002; Vampola *et al*., 2016; Yokota *et al*., 2019; Zhang *et al*., 2019). Among these, lumped-element models employing non-linear mechanical components, like coupled mass-spring-damper systems, hold particular utility for comprehensive investigations of underlying glottal phenomena (Erath *et al*., 2013). These models include biomechanical aspects of VF tissues (Titze and Story, 2002), the transversal mucosal wave (Story, 2002), aeroacoustic interactions (Horáček *et al*., 2007), glottal geometry (Galindo *et al*., 2017), and laryngeal muscle control (Titze and Hunter, 2007). A notable characteristic of lumped-element models is their ability to strike a well-suited trade-off between computation time and physiological significance.

While numerical models have traditionally been employed to simulate physiological mechanisms in modal and typical phonation, ongoing refinements have expanded their applicability to variables describing vocal function in pathological contexts. Modeling of vocal function provides a platform for exploring biomechanical aspects of impaired phonation, including the identification of biomechanical parameters and alterations in certain vocal pathologies (Jiang *et al*., 2006; Samlan and Story, 2017; Tao and Jiang, 2007), VF geometrical properties (Lucero *et al*., 2019; Mattheus and Brucker, 2011; Smith *et al*., 2013), variations in intrinsic muscular control of the larynx (Chhetri *et al*., 2009; Manríquez *et al*., 2019), and changes in the aeromechanical-acoustic interaction (Erath *et al*., 2019; Story, 2002; Zhang, 2018).

VF asymmetry holds significant relevance for clinical voice research, potentially playing a pivotal role for studying the onset of dysphonias and other vocal disorders (Pickup and Thomson, 2009). Prior studies employing modeling techniques have investigated asymmetries in VF to comprehend their underlying etiology and explore potential clinical interventions (Lucero *et al*., 2020; Mehta *et al*., 2011; Zhang and Luu, 2012). However, within the context of lumped-element models, asymmetry has primarily focused on geometrical and mechanical distinctions between the folds (Dresel *et al*., 2006; Erath *et al*., 2011; Mehta *et al*., 2011; Sommer *et al*., 2013; Steinecke and Herzel, 1995; Xue *et al*., 2010).

Motivated by the need of relating physiological muscle activation and asymmetric VF vibration, our work introduces a novel approach through the asymmetric triangular body-cover model. This model explores imbalanced activation between the left- and right-side intrinsic laryngeal musculature, providing a more comprehensive understanding of vocal asymmetry in terms of clinical descriptions (Desjardins *et al*., 2022). The proposed model adheres to the triangular shape of the glottis and the layered structure of each VF and has been integrated into a physiological vocal synthesizer, accounting for physical interactions both at the glottal level and along the vocal tract.

This work comprises distinct components, detailed in the subsequent sections. Section II delineates the adaptation of a previous model for a two-fold configuration, outlines model simulations, introduces asymmetry feature calculations, and presents two cases with high-speed camera videos for contrasting model responses with experimental data. Section III shows the results obtained through the proposed model and compares them with the asymmetries from the experimental data. Section IV offers a thorough discussion of the findings and section V concludes the manuscript.

## II. METHOD

### A. Asymmetrical Triangular Body-Cover Model

The Triangular Body-Cover Model (TBCM) (Alzamendi *et al*., 2022; Galindo *et al*., 2017), a symmetrical low-order model of the VFs, serves as the foundation for our asymmetric triangular body-cover model (a-TBCM). The TBCM incorporates tissue-fluid-acoustic interactions at the glottis, enabling simulations of sustained vowels and time-varying glottal gestures.

Building upon prior efforts in low-order modeling (Titze and Story, 2002), VF posturing (Titze and Hunter, 2007), body-cover VF models (Story and Titze, 1995), and the triangular shape of the glottis (Birkholz *et al*., 2011; Galindo *et al*., 2017), the TBCM consists of paired three-mass body-cover systems (see Figure 1 (top)) and a muscle-controlled model of all five intrinsic laryngeal muscles: cricothyroid (CT), thyroarytenoid (TA), lateral cricoarytenoid (LCA), interarytenoid (IA), and posterior cricoarytenoid (PCA).

**FIG. 1:**
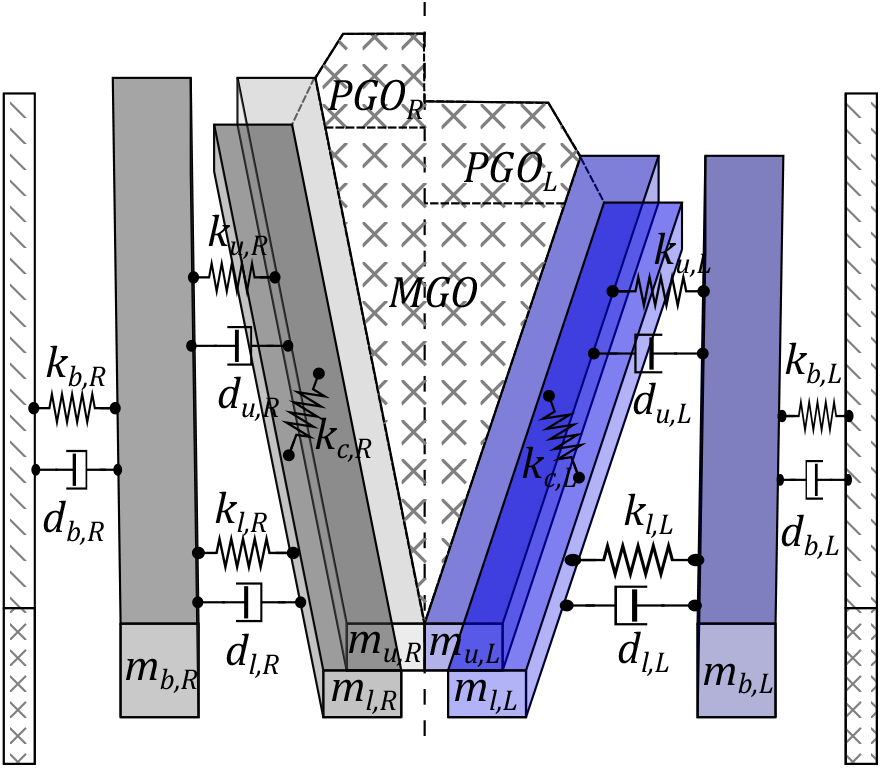
(Color online) Schematic diagram of the VF configuration according to the a-TBCM for an abducted glottal configuration.

These muscles allow for dynamic control of the prephonatory posture (i.e., vocal process accommodation) and the VF configuration (i.e., viscoelastic VF properties) during phonation. Each intrinsic muscle is treated using a modified Kelvin model for the stress-strain response in the tissues, considering both the passive and active stress components (Hunter *et al*., 2004). The model offers flexibility by allowing normalized (beween 0 and 1) individual actuation levels for each muscle, namely, (*a*_LCA_, *a*_IA_, *a*_PCA_, *a*_CT_, *a*_TA_). Moreover, this model allows for the incorporation of pulmonary pressure (*P*_*L*_) as an adjustable input parameter, working in tandem with the muscle activation profiles.

Aerodynamic forces over the VF cover layer are computed according to Titze and Story (2002), while elastic, damping, and collision forces over the body-cover masses follow Galindo *et al*. (2017). Glottal airflow is computed from the acoustic driving pressures on the glottal area, as per Lucero and Schoentgen (2015); Titze (2006); Zañartu *et al*. (2014). The acoustic wave propagation is simulated using the wave reflection analog scheme, modeling subglottal and supraglottal tracts as a discrete concatenation of acoustic cylinders with variable crosssectional areas (Zañartu, 2006).

The a-TBCM is constructed around the coupling between two independent muscle-controlled TBCMs, as illustrated in Figure 1, representing the left and right VFs. In this scheme, two muscle activation vectors are defined:

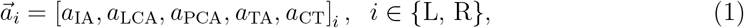

where the subscript *i* denotes the side of the a-TCBM in question, with L and R denoting left and right, respectively. Equal left/right activation vectors correspond to the symmetric control scheme in the TBCM. Then, the posture and the viscoelastic properties of each VF are defined by its own muscle activation. To model the dynamics of the VF, i.e. the movement of the body-cover elements, it is necessary to formulate and solve the coupled equations of motion for the left and right VF (see the Appendix).

### B. Muscle Imbalance for a-TBCM: Bilateral Posture and Viscoelasticity

Muscle imbalance refers to a slight difference in the activation of the laryngeal muscles between the left and right VF. These varying muscle activations serve as inputs to the control mechanism, influencing the mechanical and geometric properties of the VF. The proposed a-TBCM provides a platform to explore the impact of imbalanced activation in the intrinsic musculature on the geometric and viscoelastic properties, influencing VF oscillations and the aerodynamics of glottal airflow. To establish a reference activation condition for the intrinsic laryngeal muscles, a simplification was introduced. The activation levels of the primary adductors (LCA and IA muscles) were combined by setting *a*_LCA_ = *a*_IA_, streamlining the adjustable variables for sustained VF oscillations. Additionally, the PCA activation (*a*_PCA_) was consistently set to zero across all simulations, neutralizing its significant abductor effect on glottal posture (Alzamendi *et al*., 2022). For the TA and CT muscles, activations *a*_TA_ and *a*_CT_ were calibrated to produce voiced sounds within the fundamental frequency range of 90 to 100 Hz, aligning with the physiological characteristics of a male modal voice. The activation levels were determined based on muscle activation plots reported in Alzamendi *et al*. (2022). The reference activation for the right side is represented by:

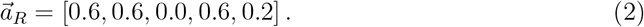

To represent the imbalance during voiced phonation, an asymmetry factor *q* ∈ [0.5, 1.5] is introduced between the left and right sides in the a-TBCM. A similar idea has been previously applied by Steinecke and Herzel (1995), henceforth SH95. However, rather than controlling asymmetries in the mass-spring elements through a gain parameter as in SH95, the a-TBCM applies the asymmetry factor directly to the muscle activation levels. These levels jointly influence the posture and configuration of the VF. For comparison with other fold asymmetry approximations, the equivalent of SH95 for the TBCM model is implemented, where:

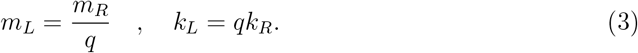

The modified masses and springs correspond to those presented in Figure 1, representing the upper, lower, and body blocks. Two primary scenarios of muscle imbalance were explored: an asymmetric activation predominantly affecting glottal adduction (vocal process) and an asymmetry in the biomechanical properties (mass-spring values) of VF. To represent the first case, the asymmetry factor *q* was applied to *a*_LCA_ and *a*_IA_ in the left VF. The muscle activation vector for the left VF is then given by:

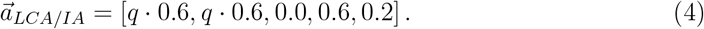

For asymmetries in the biomechanical properties, *q* was applied to *a*_CT_ and *a*_TA_ simultaneously, considering the CT-TA antagonistic relationship in determining the viscoelastic (VE) properties in the VF (Titze and Story, 2002). The muscle activation vector for the left VF in this scenario is expressed as:

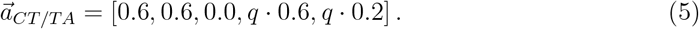

Additionally, individual muscle imbalance in each CT and TA was studied separately, given its significance in characterizing the glottal configuration in the model, based on the work of Titze and Hunter (2007); Titze and Story (2002). This resulted in a total of four configurations of muscular imbalance and the direct imbalance in masses and springs The simulations were performed with a driving pulmonary pressure of 1 kPa. A truncated Taylor-series approximation was implemented to simultaneously solve the differential equations of motion for the six masses, using a sampling frequency of 44.1 kHz. The glottal tract was defined as the area function corresponding to an /*i*/ vowel (Story, 2008), and a subglottal tract as an inverted cone shape (Zañartu, 2010; Zañartu *et al*., 2014).

### C. Model Derived Measures

Several measures were derived from a-TBCM simulations to establish differences between the muscle imbalance scenarios, study the effect of the asymmetric factor on VF vibratory asymmetries, and compare the model with certain clinical cases. These measures include left-right amplitude asymmetry (AA), left-right phase asymmetry (PA), fundamental frequency (*f*_*o*_), and open quotient (OQ). Following Mehta *et al*. (2011), synthetic kymograms were generated from the right and left VF edge positions over time as illustrated in Figure 2. Based on these representations, measures of the VF oscillatory asymmetries can be defined:

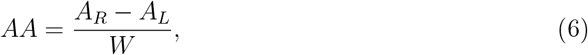

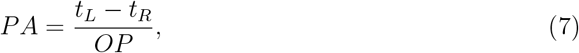

**FIG. 2:**
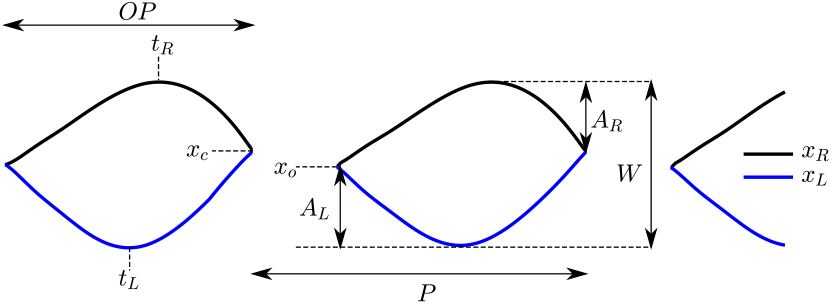
(Color online) Kymogram parametrization: *x*_*L*_, *x*_*R*_ displacement of left/right VF; *t*_*L*_, *t*_*R*_ time of maximal left/right VF displacement; *x*_*c*_, *x*_*o*_ mediolateral position of VFs at the glottal closure/opening; *A*_*L*_, *A*_*R*_ maximum displacement of left/right VF; *W* peak-to-peak glottal width; *P* glottal period; and *OP* open phase. Adapted from (Mehta *et al*., 2011)

Note that the normalization factor *OP* for defining PA follows that of Mehta *et al*. (2011), where the focus is on the acoustic effects generated during the open phase. These measures assess different aspects of oscillatory asymmetries. The left-right amplitude asymmetry, Equation 6 (Qiu *et al*., 2003), is the ratio between the difference and the sum of the maximum excursions for the right (reference) and left (affected) VF amplitude traces within one vibration period. The left-right phase asymmetry, Equation 7 (Bonilha *et al*., 2008; Lohscheller *et al*., 2008), is the time delay between the maximum excursions of the left and right VF normalized by the oscillation period.

The open quotient is also computed from the kymogram as the ratio of the duration of the open phase (OP) to the period (P) for each cycle.

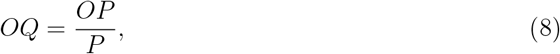

### D. Clinical Cases

In conjunction with the modeled scenarios, this study draws support from in vivo high-speed video (HSV) recordings representing voice disorders. Two participants, one with non-phonotraumatic vocal hyperfunction (muscle tension dysphonia) and another with unilateral VF paralysis, were included. During data acquisition, participants produced a sustained /*i*/ vowel sound at optimal pitch and loudness levels, adhering to informed consent procedures and Institutional Review Board (IRB) protocols outlined in FONDECYT 1151077 project. The HSV data collection involved an advanced high-speed camera (SA-X2, Photron, Tokyo, Japan) synchronized with a rigid endoscope (9106, KayPentax, Montvale, NJ) equipped with a 35 mm C-mount adapter and a xenon light source (7152B, KayPentax, Montvale, NJ). The original HSV dataset, comprising approximately 2,670 frames at 8,000 frames per second (fps), underwent pre-processing. This included selecting 600 frames and identifying a 256 × 256 pixel region of interest containing only the VFs. The refined HSV dataset was then used to segment the glottis area using GlottalImageExplorer (Birkholz, 2016).

Subsequent to the glottis area segmentation, the digital kymogram (DKG) technique proposed by Mehta *et al*. (2011) was applied to visualize VF trajectories. These lateral displacement waveforms, capturing intricate motions of the left and right VFs, were traced along a one-pixel line positioned at the midpoint of the glottal region’s posterior-anterior length. Additionally, the DKG outputs facilitated the automated extraction of amplitude and phase asymmetry metrics for the left and right VFs.

## III. RESULTS

### A. Influence of Muscle Imbalance on VF Properties

As shown in Figure 3, the asymmetry in the adductor muscles (LCA/IA) has a negligible impact on the mass of the blocks but leads to a minor alteration (<10%) in the elastic component. This is due to the relatively minor role that adductor muscles play in the construction of the folds, although they do contribute slightly to their elongation.

**FIG. 3:**
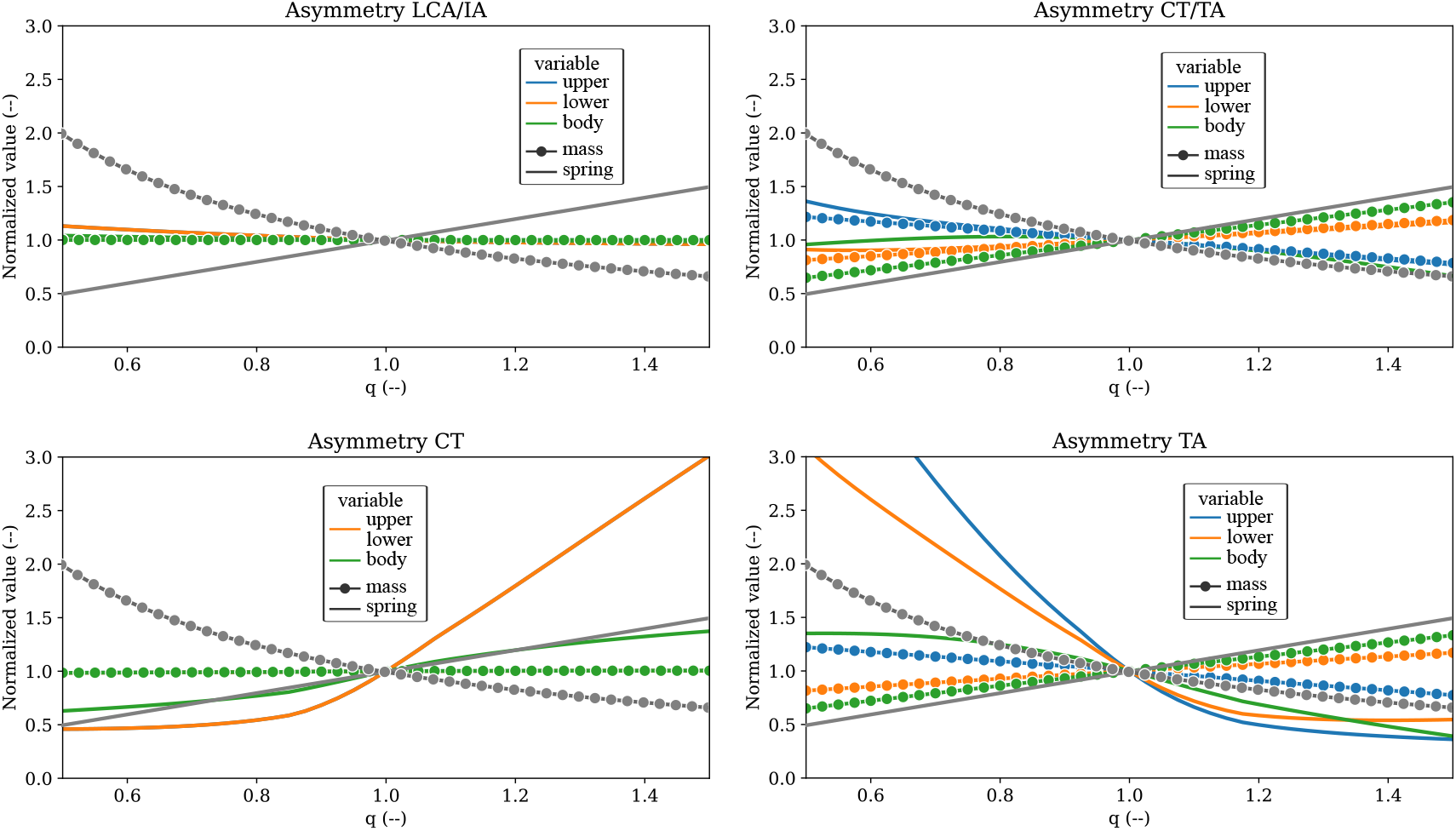
(Color online) Effects on normalized masses (pointed line) and springs (solid line) when varying the joint asymmetry factor *q* for VP (top left), VE (top right) configurations, and single asymmetry in CT (bottom left) and TA muscle (bottom left). The legend represents the nomenclature in a-TBCM scheme (see Figure 1) form mass and spring: upper, lower, and body blocks. The gray lines describes the effects due to the asymmetry mechanism introduced in SH95.

The imbalance between CT and TA produces opposing effects on the elastic component of the VF. Imbalance in the CT muscle generates a trend in the same direction as the SH95 reference mechanism, though not in a strictly linear manner. Conversely, an imbalance in the TA muscle produces an effect that varies from positive to negative as the value of *q* increases, opposite to the reference mechanism.

Regarding the mass component, an imbalance in the CT muscle does not lead to any changes. However, an imbalance in the TA muscle results in a linear change in the mass of the body block and redistributes the mass of the cover between the upper and lower blocks.

This pattern contradicts the direct imbalance in mass and spring values and highlights the intricacy of the laryngeal control mechanism. The CT muscle elongates the fold, impacting its elasticity without adding any mass. In contrast, the TA muscle is responsible for adding mass to the fold and redistributing it between the blocks by altering their dimensions. This generates an effect on both the mass and spring components.

Figure 4 illustrates the impact of *q* on fundamental frequency (*f*_*o*_) and posterior glottal opening (PGO) in various imbalance scenarios. Notably, the configuration with an imbalance in LCA and IA muscles exhibits a non-zero posterior gap, resulting in an increased *PGO* for *q* < 1. This effect arises from the action of the LCA and IA muscles in adducting the VFs. In contrast, other imbalance scenarios do not induce changes in PGO due to a lack of alteration in the adduction degree of the folds. For instance, the SH95 approach, which does not consider vocal posture, exhibits no *PGO* changes.

**FIG. 4:**
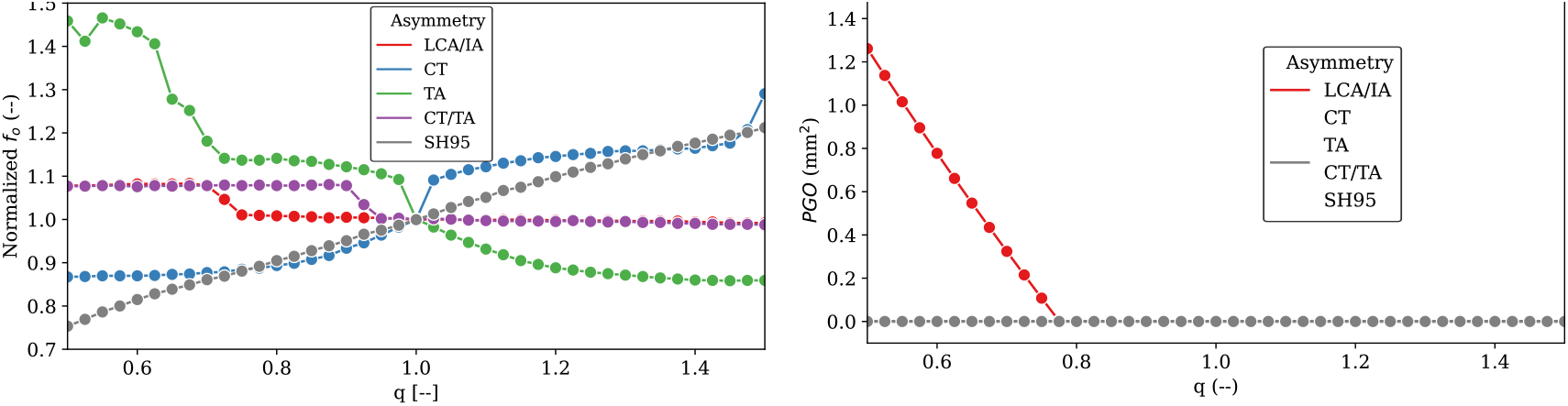
(Color online) Effects on *f*_*o*_ and *PGO* when varying *q* for muscle imbalance configurations. SH95 results shown for comparison.

Upon examining the influence of the *q* factor on fundamental frequency, distinct behaviors emerge in comparison to the SH95 model. In the SH95 model, the *q* factor proportionally affects *f*_*o*_, reflecting its design intent. Conversely, in cases of muscle imbalance, different patterns are observed. An imbalance in the TA muscle leads to an inverse relationship with *f*_*o*_—increasing *q* results in a decrease in *f*_*o*_, influenced by the effect of TA on the VF spring. On the other hand, an imbalance in CT exhibits a slight direct correlation with *f*_*o*_; an increase in q correlates with an increase in *f*_*o*_, with a defined threshold for the minimum and maximum *f*_*o*_ values. In scenarios involving joint imbalance of two muscles, CT/TA and LCA/IA, the fundamental frequency displays a step-like behavior. At a specific point, there is a 10% variation in its value, indicating a transition between frequencies.

### B. Influence of Muscle Imbalance on VF Oscillation

To examine the impact of muscle imbalance on VF oscillation, Figure 5 provides waveforms illustrating synthetic kymograms and glottal airflow. These visualizations encompass the baseline a-TBCM simulation and imbalance scenarios under two configurations: hypo-function (q = 0.5) and hyperfunction (q = 1.5). The first row displays a synthetic kymograph illustrating the amplitude displacements of the left and right VFs in millimeters. The second row presents glottal airflow expressed in milliliters per second. All signals correspond to a 50 ms simulation. Additionally, Table I furnishes computed asymmetry measures, including *f*_*o*_, *OQ, AA*, and *PA* for the simulated scenarios.

**TABLE 1:** Comparison of asymmetry values and oscillation features between the baseline and the q configurations shown in Figure 5. For each feature, the first row is for the hypofunction (q = 0.5) and the second row is for the hyperfunction (q = 1.5)

|  | Baseline | SH95 | LCA/IA | CT | TA | CT/TA |
| --- | --- | --- | --- | --- | --- | --- |
| $f_o$ (Hz) | 96 | 72.4 | 103.6 | 83.4 | 140.2 | 103.5 |
|  |  | 116.5 | 95.3 | 124 | 82.6 | 94.5 |
| $OQ$ (%) | 49 | 60 | 66.5 | 51.8 | 78.7 | 68.4 |
|  |  | 53.9 | 49.1 | 72.8 | 52 | 49.2 |
| $AA$ (%) | 0 | 31.3 | -1.5 | -3.1 | -21.5 | -8.27 |
|  |  | -22.1 | 0.4 | -8.7 | -0.5 | 2.9 |
| $PA$ (%) | 0 | 24.71 | -1.9 | 18.2 | -17.5 | -5.8 |
|  |  | -21.5 | 0.44 | -16.9 | 18.2 | 0.3 |

**FIG. 5:**
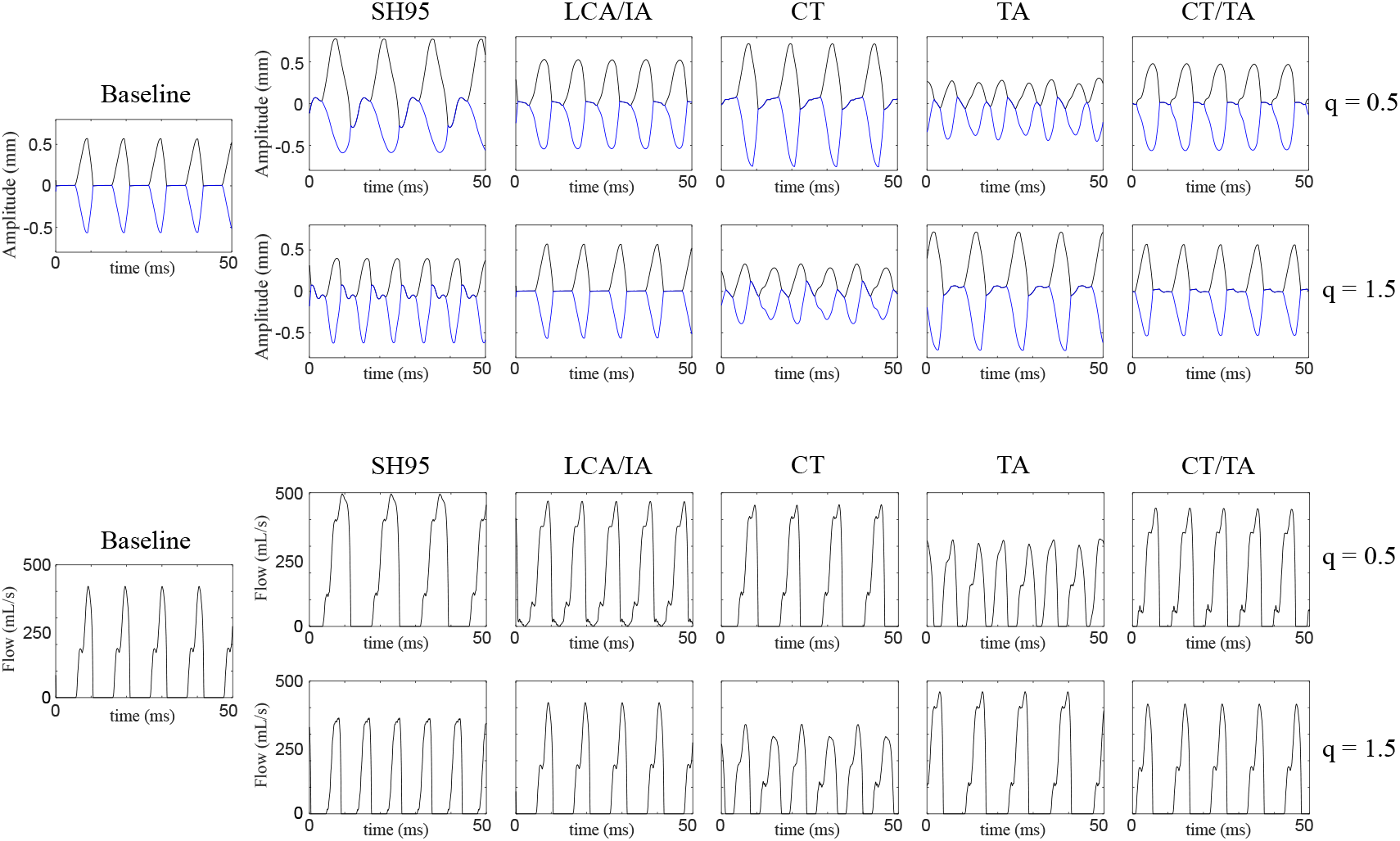
(Color online) Comparison of kymograms and airflows from a-TBCM simulations for asymmetry scenarios and two imbalance conditions: hypofunction (*q* = 0.5) and hyperfunction (*q* = 1.5.)

In the context of direct mass and spring imbalance, as depicted in the second column of Figure 5, the hypofunction configuration results in an augmented glottal area, indicated by an increase in airflow. Furthermore, the oscillation amplitude of the affected fold is smaller compared to the unaffected fold, leading to an *AA* value exceeding 30%. A positive *PA* value, surpassing 20%, is also evident, accompanied by a decrease in the fundamental frequency during oscillation. In the hyperfunction configuration, this displacement behavior is reversed. Now, the affected fold undergoes greater displacement, reaching its maximum value first, accompanied by an increase in frequency and a decrease in flow amplitude.

In the case of imbalance in the LCA/IA adductor muscles, hypofunction exhibits a slight increase in maximum airflow, along with asymmetry measurements *AA* and *PA* below 2%. These measurements indicate that the less adducted (modified) fold exhibits greater displacement. A notable observation in this case is the nonzero airflow value during the closed phase of VF oscillation, aligning with the nonzero *PGO* value reported in Figure 4. In the hyperfunctional configuration, this asymmetry maintains the baseline shape, as, after a certain level of activation in LCA and IA, the folds are already parallel. Therefore, an increase in muscle activation does not induce visible changes in the glottis with asymmetry measures below 1%.

For individually applied CT and TA muscle imbalance, a cross-behavior is observed in terms of hypo- and hyperfunction. In the case of altered CT muscle activity, displacements comparable to the baseline are observed in hypofunction, with greater displacement of the affected fold (*AA*=-3%) and reduced amplitude oscillation for the hyperfunctional condition. Conversely, in TA imbalance, the reduced amplitude of the folds occurs in hypofunction, while oscillation similar to the baseline is observed in hyperfunction. This underscores the antagonistic nature of this muscle pair in the viscoelastic composition of the VFs, as also observed in the mass and spring curves presented in Figure 3. This cross-behavior is clearly evident in the *OQ, f*_*o*_, and *PA* values presented in Table I. For individual muscle imbalance, positive values for AA are unattainable, signifying that the affected muscle has a greater amplitude of oscillation.

In terms of VF oscillation behavior, Figure 7 demonstrates the influence of the asymmetry factor *q* on *AA* and *PA* across the examined cases. The conventional imbalance scheme, SH95, yields positive values for *AA* and *PA* when *q* < 1, indicating that the affected fold’s oscillation is dampened due to an increase in mass and a decrease in elasticity. Conversely, it intensifies its oscillation and frequency, resulting in negative values of AA and PA for q > 1.

**FIG. 6:**
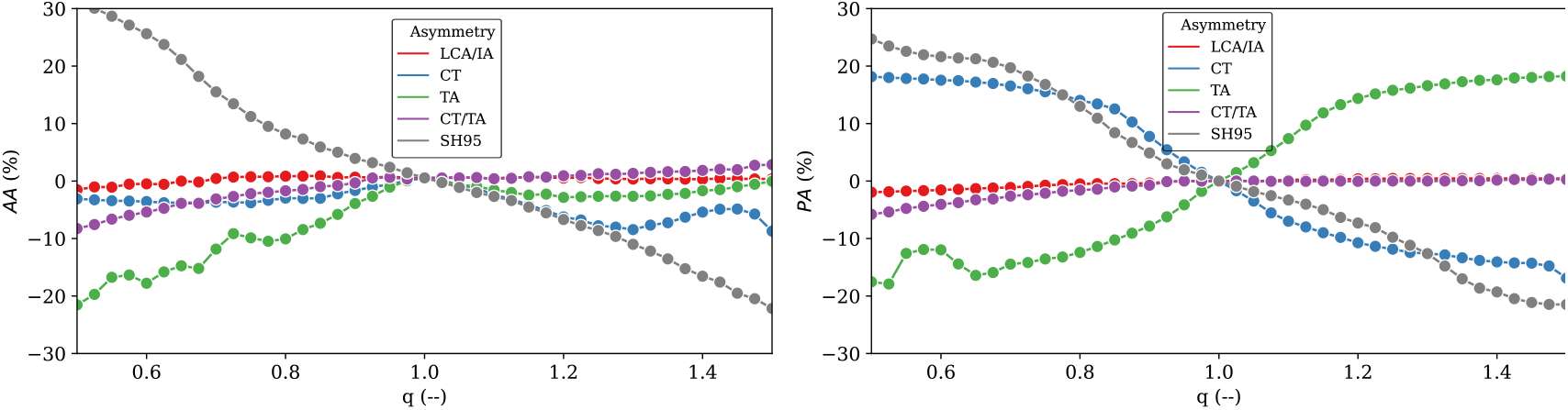
(Color online) Effect of changing *q* on asymmetry metrics; (Left) *AA* and (Right) *PA*, for the different imbalance approaches.

**FIG. 7:**
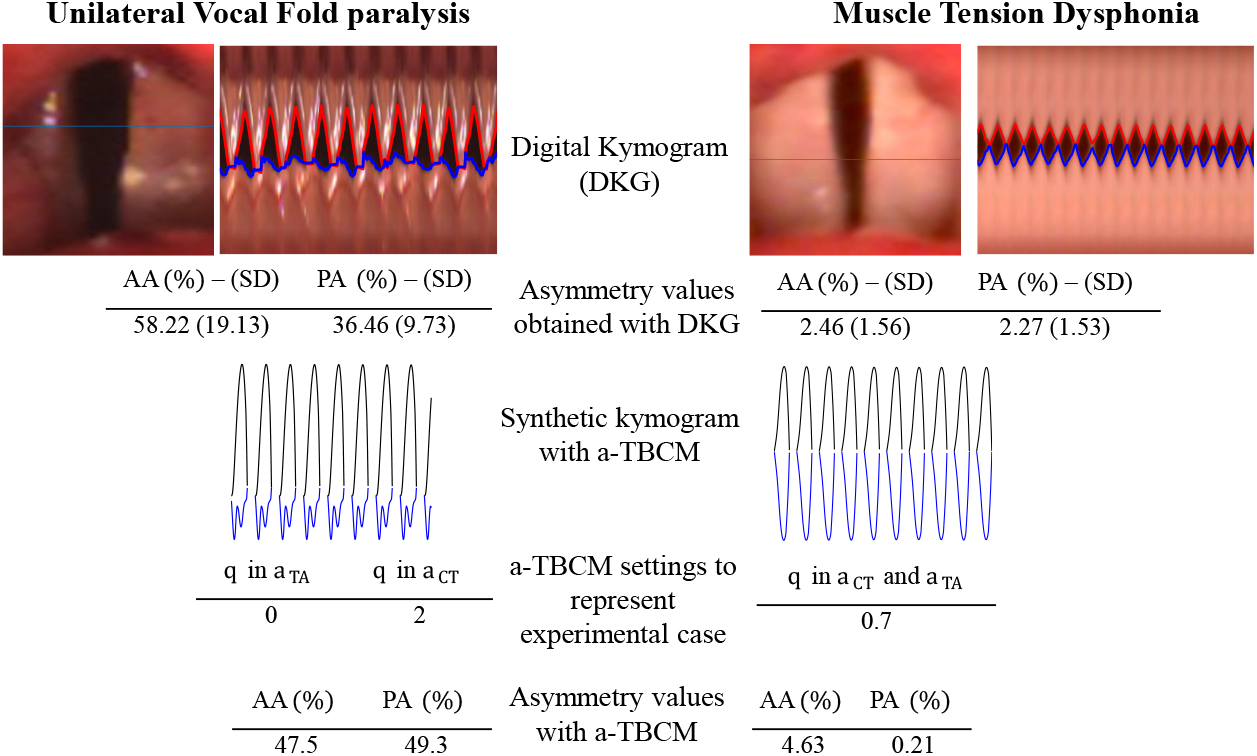
(Color online) Comparison of clínical (HSV and DKG) data and the corresponding simulated responses using a-TBCM: (left) Unilateral VF paralysis and (right) Muscle Tension Dysphonia.

When considering scenarios of muscle imbalance, several significant observations emerge. The imbalance in CT mirrors the conventional scheme’s trend for *PA*, while the imbalance in TA exhibits the opposite trend. This discrepancy can be attributed to the behavior of the elastic component as a function of the *q* factor, increasing in the CT imbalance and SH95, and decreasing in the TA imbalance. Regarding *AA*, it proved challenging to find a muscle imbalance scenario that generated the same range as the reference scheme. This stems from the mechanism of muscle activation control and the mass and spring parameters, where not all mass values move in the same direction when varying muscle activation.

In contrast, the imbalance in LCA/IA yields the most minor effect on asymmetry mea-surements within the studied range, resulting in asymmetries of less than 2%. Meanwhile, the combined imbalance of CT/TA amalgamates the effects of individual imbalance, producing phase asymmetries of approximately 6% and amplitude asymmetries approaching 10%.

### C. Contrasting a-TBCM Against In Vivo Examples

To demonstrate the efficacy of incorporating muscle imbalance in modeling various clinical scenarios, we present clinical HSV images alongside simulated data generated using the a-TBCM in Figure 7. This illustration highlights two distinct cases: unilateral VF paralysis (left) and muscle tension dysphonia (right). Each case features the DKG derived from clinical data, juxtaposed with the synthetic kymogram produced by the a-TBCM. Synthetic DKGs were generated by adjusting amplitude and phase asymmetry measurements heuristically to align with their respective clinical mean and variance values, employing the muscle activation defined in Equation 5 as the baseline. The simulation process encompasses the insights detailed in the preceding subsections, accounting for the characteristic oscillatory behavior associated with each pathology. Specifically, for VF paralysis, our aim was to emulate a rigid VF, while for muscle tension dysphonia, we introduced a subtle differentiation between the VFs.

Figure 7 highlights the specific imbalance factors employed in generating the synthetic kymogram, providing clarity on the value of q and the muscle to which it is applied. It is worth noting that while this approach may not be entirely singular, a more precise representation could potentially be achieved through model optimization considering the complete kymogram signal. Nevertheless, in both cases, the a-TBCM successfully produces a waveform closely mirroring the actual kymogram. In the instance of VF paralysis, this results in a fold with minimal amplitude oscillation, out of phase in time. In the case of muscle tension dysphonia, it yields a slightly asymmetric oscillation in both amplitude and phase.

Figure 7 also presents corresponding *AA* and *PA* measurements from both clinical and synthetic data. Clinical assessments of VF oscillatory asymmetry display noteworthy variability, emphasizing their sensitivity to spatiotemporal resolution and the challenges in obtaining precise measurements of this nature. In the case of unilateral paralysis, both *AA* and *PA* values surpass 30%, indicating substantial asymmetry. These asymmetry values are effectively captured by the a-TBCM, which deactivates the TA in the paralyzed fold. Similarly, in the case of muscle tension dysphonia, the a-TBCM enables the representation of such levels of asymmetry by introducing an activation imbalance in the CT and TA simultaneously.

## IV. DISCUSSION

The primary aim of this study was to introduce and demonstrate the concept of muscle imbalance as a mechanism for inducing asymmetries within the VFs. We sought to establish its potential in characterizing clinical scenarios by examining the influence of intrinsic laryngeal muscles on the glottis and drawing comparisons with previous approaches for understanding asymmetric VF oscillations.

Our findings lead us to conclude that achieving a direct differentiation scheme akin to SH95 through an imbalance in a single muscle alone is not feasible. The closest approximation is attained through an imbalance in the CT muscle. Additionally, the antagonistic relationship between the CT and TA muscles hinders the replication of the direct imbalance scheme. However, introducing imbalance in both these muscles concurrently, albeit in opposite directions with proportional imbalance in CT and inversely proportional in TA, appears to be a promising avenue towards achieving an analog of the conventional asymmetry mechanism using muscle imbalance. It is worth noting that a comprehensive exploration of this intricacy exceeds the scope of this initial study.

The proposed method for prescribing muscle imbalance provides a systematic means to introduce asymmetry in the composition and positioning of the VFs. This asymmetry has been studied independently by other researchers. Prior efforts (Dresel *et al*., 2006; Jiang *et al*., 2006; Mehta *et al*., 2011; Xue *et al*., 2010) report ranges of amplitude and base asymmetry akin to our findings. Conversely, studies such as (Dresel *et al*., 2006; Samlan and Story, 2017; Samlan *et al*., 2014) report differentiation in the vocal process without altering the mechanical properties of the VFs, resulting in increased minimum glottal flow and decreased sound pressure, much like the effects observed with the imbalance in the LCA and IA adductor muscles.

When considering the results of mechanical parameters and asymmetry measurements collectively, it becomes evident that phase asymmetry aligns with the spring component trends, while mass trends are associated with amplitude asymmetry. This elucidates why the imbalance mechanism does not yield significant amplitude values. Each block follows a distinct mass pattern when subjected to imbalance, resulting in a small AA that deviates from the SH95 scheme, where all blocks experience similar mass effects.

The clinical examples presented in this study serve to illustrate the potential of our approach in emulating actual cases, thereby enhancing our understanding of the origins of differences in VFs, be it in cases of vocal cord paralysis (Ivey, 2019; Tipton *et al*., 2020) and primary muscle tension dysphonia (Hsiung and Hsiao, 2004; Spencer, 2015). While similar insights may exist within other methodologies, our approach integrates muscular activity directly, impacting glottal configuration.

For instance, paralysis is depicted as an inert and minimally mobile fold, whereas further variations showcase a fold with high mass and rigidity. Notably, this work represents an initial foray into a parameter fitting and optimization problem, without delving into extensive details. Our objective is to demonstrate the utility of this imbalance approach, as exemplified in Figure 7.

However, studying VF oscillation with diverse properties presents several challenges, including the presence of harmonics, as illustrated in Figure 5, a topic extensively explored by other researchers (Sommer *et al*., 2013; Steinecke and Herzel, 1995; Zhang and Jiang, 2004). Another critical consideration is the impact of the chosen reference point, whether it pertains to muscular activation or the specific values assigned to masses and springs. Therefore, future endeavors should encompass these aspects, in conjunction with the integration of clinical expertise, to ensure the accuracy of predictions and trends generated by models representing asymmetric VF oscillations.

## V. CONCLUSION

This study introduces a muscle-controlled, triangular and asymmetrical body-cover model of the VFs to explore the impact of muscle imbalance on VF vibration. The findings reveal significant alterations in the dynamics of mass-spring and modulation of fundamental frequency when comparing activation-based muscle imbalance to conventional methods. The incorporation of muscle activation as a distinguishing factor in VF composition has the potential to revolutionize our comprehension of vocal asymmetry. This shifts the focus beyond mere structural considerations, emphasizing the critical role of muscular coordination. Moreover, initial experiments have successfully replicated clinical scenarios of VF paralysis and muscle tension dysphonia. However, further research is imperative to comprehensively assess the proposed muscular imbalance descriptions in clinical settings. Nevertheless, this study makes a significant contribution to advancing our understanding of VF dynamics and highlighting the pivotal role of muscle imbalance in normal and disordered voice production.

## ACKNOWLEDGMENTS

This research was supported by the National Institutes of Health (NIH) National Institute on Deafness and Other Communication Disorders grant P50 DC015446, ANID grants Becas de Doctorado Nacional 21202490 and 21190074, FONDECYT 1230828, BASAL FB0008, UTFSM grant DPP PIIC N∘ 077/2022, and Corporación Universitaria Iberoamericana, Research Department, under search grant for project 202210D038. The content is solely the authors’ responsibility and does not necessarily represent the official views of the National Institutes of Health.

## APPENDIX: IMPLEMENTATION OF THE ASYMMETRICAL TRIANGULAR BODY-COVER MODEL

The proposed a-TBCM is, an extension of the TBCM (Alzamendi *et al*., 2022; Galindo *et al*., 2017), builds upon prior efforts (Birkholz *et al*., 2011; Galindo *et al*., 2017; Story and Titze, 1995; Titze and Hunter, 2007; Titze and Story, 2002). The proposal moves from a symmetrical to an asymmetrical glottal configuration considering the left and right VFs with two VFs with their independent muscle activation vector.

### A. Posture and Biomechanical properties

For each side of the a-TBCM, as shown in Figure 1, the dynamic adjustment of vocal posture and VF configuration as a function of activation vector 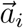 in Equation 1 follows the methodology in the TBCM. The theoretical development and the implementations were described in Alzamendi *et al*. (2022); Galindo *et al*. (2017), and are briefly summarized for completeness.

A system of equations of motion models the laryngeal posturing by describing the relative movements between the arytenoid cartilage and the cricoid cartilage, and between the cricoid cartilage and the thyroid cartilage, in response to the forces in the laryngeal tissues. Vocal process Cartesian coordinates are tracked, and from this information and the forces in the laryngeal tissues, the adductory displacements, Δx_*u,i*_, Δx_*l,i*_, and the VF lengths, L_*i*_, for *i* ∈ {L, R} are obtained (see Figure 8).

**FIG. 8:**
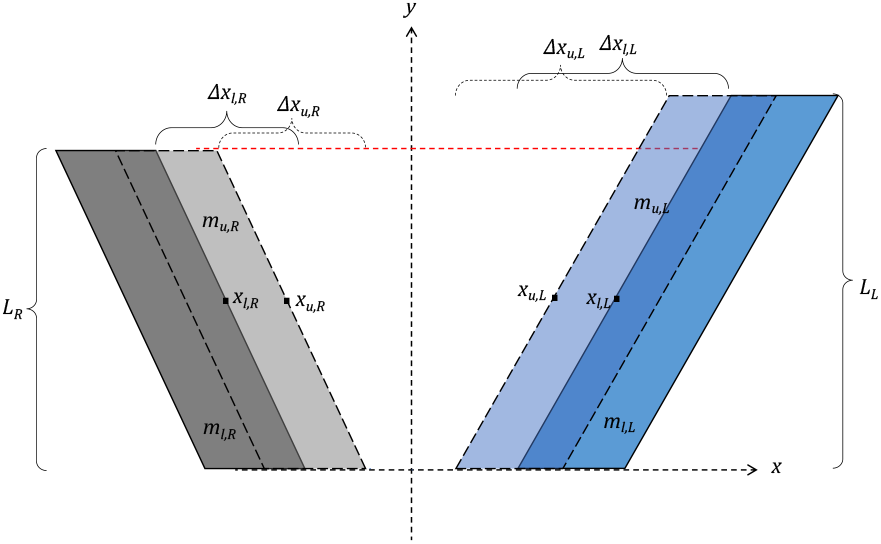
(Color online) Top view: 2D diagram describing the abducted VFs positioning for the cover blocks in the a-TBCM.

Thereupon, empirical rules are applied for adjusting the geometrical and biomechanical parameters (Alzamendi *et al*., 2022; Titze and Story, 2002) (e.g., thickness, depth, and mass m for each block, the nodal point, the glottal convergence, and the values for the spring k and damping d parameters (see Figure 1)) of the left/right TBCM.

### B. Glottal areas calculation

The total glottal area *A*_*g*_ is the contribution of the membrane portion of the VFs (MGO) and the posterior gap (PGO), as shown in the Figure 1.

In the a-TBCM, the MGO for upper or lower blocks is computed as follows:

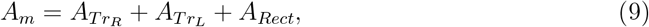

where the subscripts indicate whether the area is triangular, T*r*, or rectangular, Rect. For the computation of these areas, the fraction of the block that is under collision is introduced as a quantity that simplifies the expressions (Birkholz *et al*., 2011; Galindo *et al*., 2017):

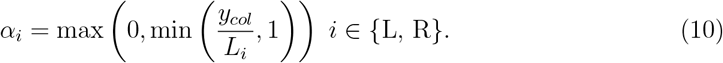

The collision height y_*col*_ in Equation 10 denotes the y-coordinate of the point where the left- and right-edge lines intercept determines the collision height (see the red dot in Figure 9). There are two possible scenarios for the glottis, as shown in Figure 9. (top) VF in no collision: (α = 0), and (bottom) VFs in collision: (α > 0).

**FIG. 9:**
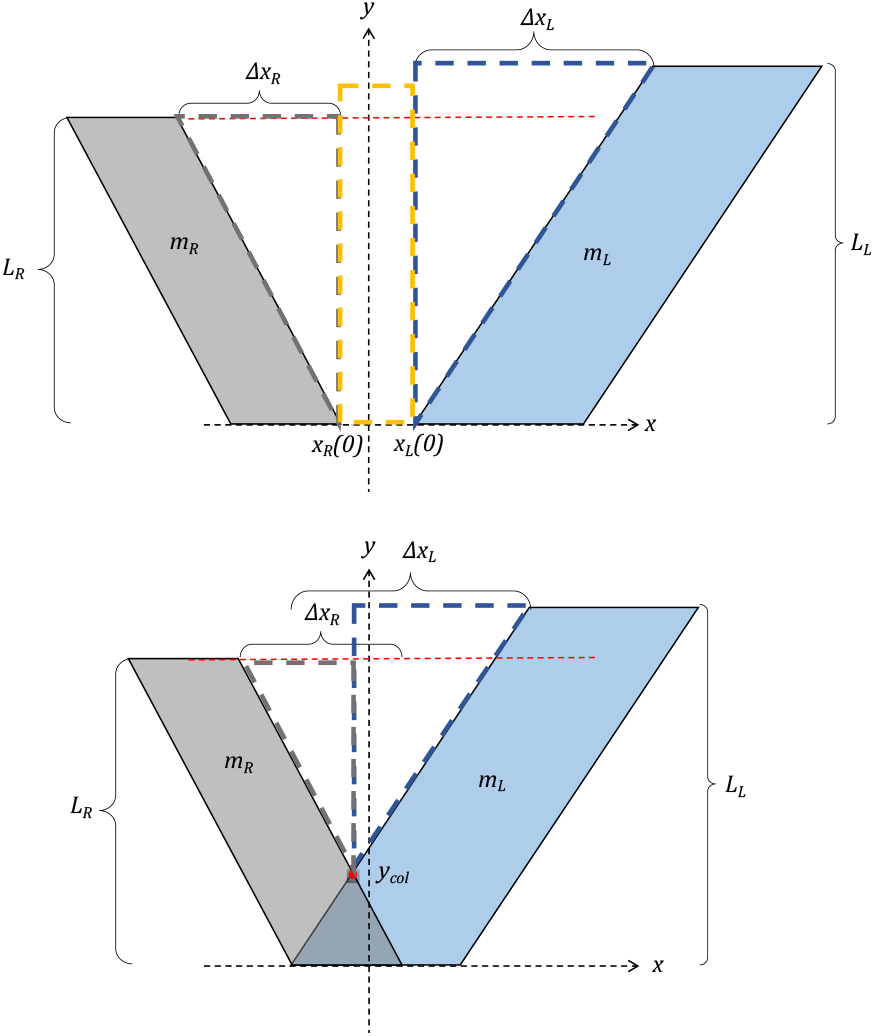
(Color online) Glottal area scheme for VF blocks: (top) no collision, (bottom) collision.

The triangular-shaped glottal area is calculated as follows:

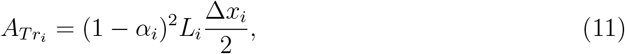

where factor (1 − α_*i*_) is the non-collision portion of the VF, and the subscript i ∈ {L, R}. On the other hand, the rectangular-shaped area is obtained from:

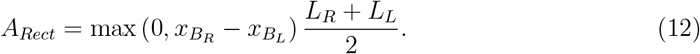

Note that during VF collision, A_*Rect*_ = 0 in Eq. (9).

The cross-sectional area, A_*t*_, and the contact area, A_*c*_, are obtained from the collision fraction, α_*i*_, in a similar way as in Alzamendi *et al*. (2022); Galindo *et al*. (2017):

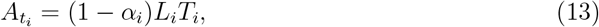

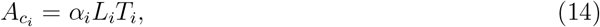

where *i* ∈ {L, R}.

The posterior glottal opening, PGO, is defined for each VF following Alzamendi *et al*. (2022), using the cricoarytenoid junction coordinate. In the a-TBCM, the PGO is the contribution of both VFs:

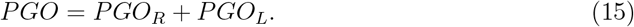

The total glottal area is the sum of the membranous area and the posterior gap:

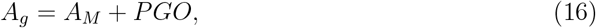

Where 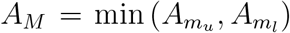, the minimum between upper and lower MGO, as shown in Figure 8.

#### C. Equations of Motion

The asymmetrical VF vibrations are simulated on the basis of coupled left/right systems of equations of motion. Each system simulates the medial-lateral displacements for the upper (*u*) and lower (*l*) cover masses and the body (*b*) mass in the TBCM (see Figure 1).

The equations of motion for each VF are:

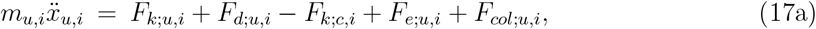

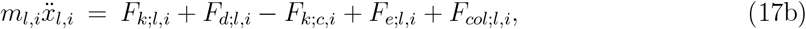

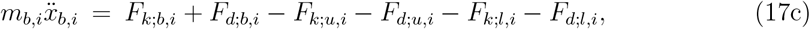

where *i* denotes the considered side (L or R), m is the mass of the block and *x* is the medial-lateral displacement over time. The right side of the equation presents the net force acting upon the block due to elastic (*k*), damping (*d*), aerodynamic (*e*), and collision (col) components.

The elastic forces (*F*_*k*;*u,i*_, *F*_*k*;*l,i*_, *F*_*k*;*c,i*_, *F*_*k*;*b,i*_) are modeled through a nonlinear Hooke’s law, and the damping forces (*F*_*d*;*u,i*_, *F*_*d*;*l,i*_, *F*_*d*;*b,i*_) are modeled proportional to the velocity. These force components are not affected by the presence of the other VF; the rules for calculation are the same as in Galindo *et al*. (2017); Story and Titze (1995). The explicit equation for each force component can be found in the appendix in Galindo *et al*. (2017).

The aerodynamic driving forces 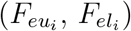 represents the force that the intraglottal pressure exerts on the VFs. It depends on the glottal flow, glottal and transversal area. Previous works explain how to calculate the intraglottal flow and pressure from the upper/lower glottal area (Story, 2008). Using the definition of pressure, the aerodynamic force for the upper/lower block is:

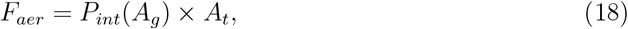

where P_*int*_ is the intraglottal pressure (constant in the entire upper or lower block) which is a function of glottal area A_*g*_ and the sub- and supra-glottal pressures, and A_*t*_ is the cross-sectional area that multiplies the pressure to compute the force component, which depends on the geometry and collision fraction α of the VF. The equations from (Galindo *et al*., 2017; Lucero and Koenig, 2005) are used to calculate the intraglottal pressure from the glottal area; however, it is necessary to define how to calculate the glottal and cross-sectional areas in the a-TBCM due to the lack of mirror symmetry.

The collision forces (F_*col*;*u,i*_, F_*col*;*l,i*_) depend on the interpenetration distances in the upper/lower cover masses due to the impact between opposing VFs. Therefore, the computation of the collision force requires defining when the collision occurs, and the VF section undergoing the collision.

Similar to Galindo *et al*. (2017), the total collision force for the lower or upper blocks is:

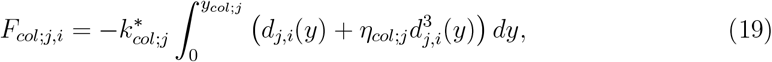

where *i* ∈ {L, R}, *j* ∈ {l, u}, *d*(*y*) is the interblock penetration distance, y_*col*_ is the collision height, 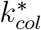 is the effective spring collision constant, and η_*col*_ is a nonlinear coefficient. To calculate the interblock penetration distance *d*(*y*), the VF edge is described as a straight line. The x position of the VF is a function of the *y* coordinate, as follows:

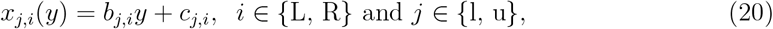

where *b* is the slope and *c* is the intercept, see below. The explicit values for quantities are obtained from Figure 8 that shows a 2D top view of the cover masses, where L_*i*_ is the VF length, *x*_*j,i*_ is the mass position and Δ*x*_*i,j*_ is posterior displacement, given the degree of abduction by the muscle activation.

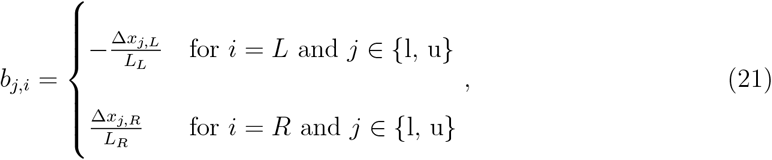

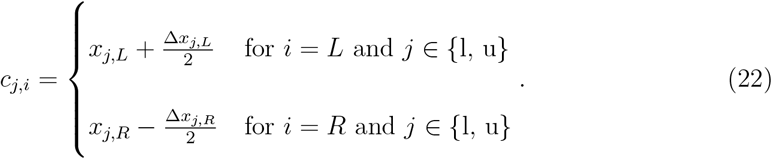

Without loss of generality, consider one of the cover elements, either upper or lower for both VFs; the upper or lower subscript is removed to have a short notation since the expression and the formulation is equivalent for both blocks. With the mathematical description of the VF edges in Equation 20, the interblock penetration distance is defined by:

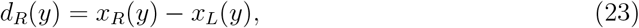

note that *d*_*L*_(*y*) = −*d*_*R*_(*y*), this denotes the opposite direction in collision forces. Additionally, this distance gives the collision condition:

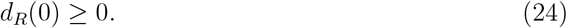

For the calculation of the collision height y_*col*_ in Equation 19, three possible glottal configurations are considered (see Figure 10): the posterior portions in both VFs are abducted (Δ*x*_R_, Δ*x*_L_ > 0), the right VF is adducted and the left one is medialized (Δ*x*_R_ > 0 and Δ*x*_L_ = 0) and *vice versa*, and both VFs are tightly adducted (Δ*x*_R_, Δ*x*_L_ = 0).

**FIG. 10:**
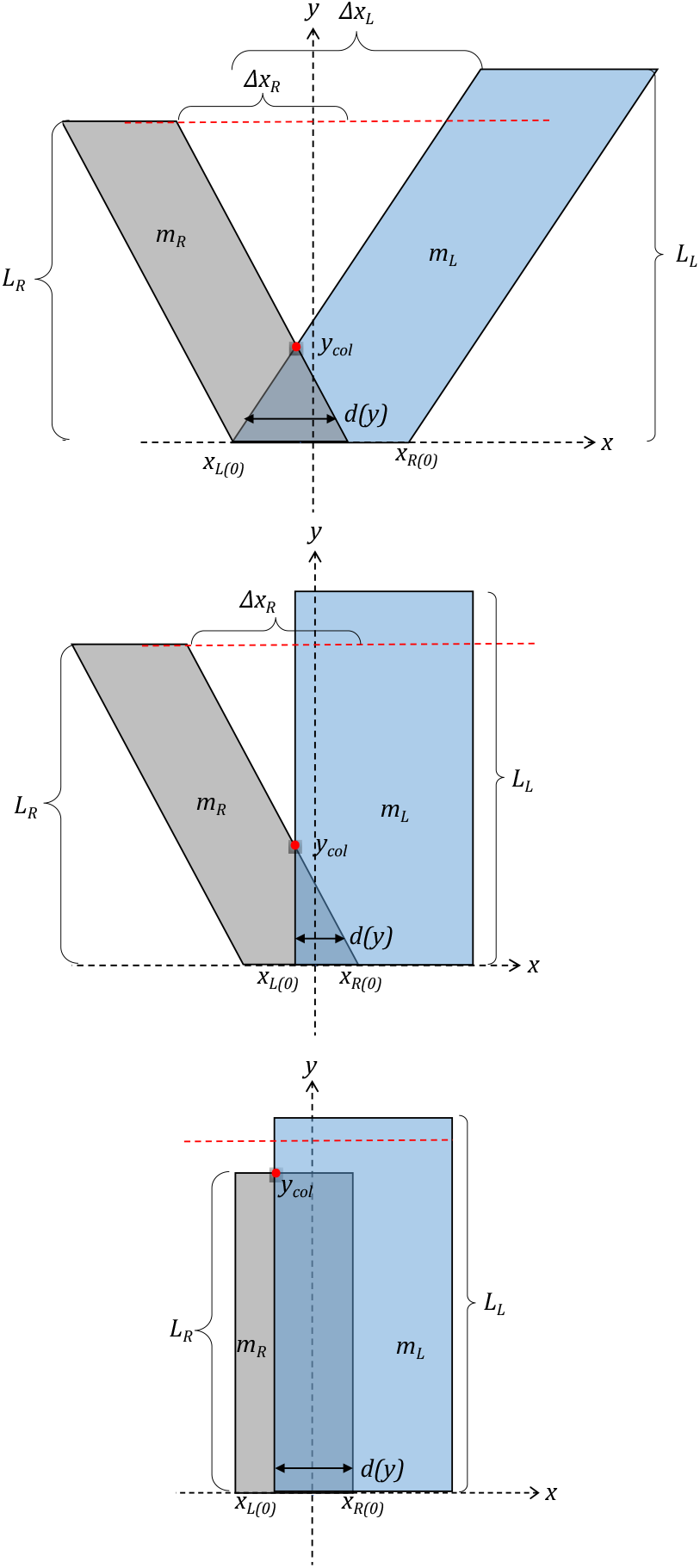
(Color online) The three collision scenarios in the a-TBCM determined by the left/right posterior displacements: (Top) case Δ*x*_R_, Δ*x*_L_ *>* 0, (Middle) Δ*x*_R_ *>* 0 and Δ*x*_L_ = 0 (and *vice versa*), and (Bottom) Δ*x*_R_, Δ*x*_L_ = 0.

For the cases where at least one of the VFs has an adduction degree:(Δ*x*_*i*_ *≠* 0), the collision height is calculated from the interpenetration distance condition:

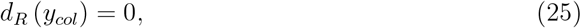

with some algebra:

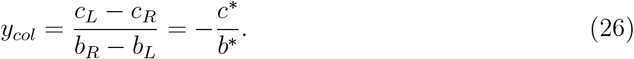

For the case of parallel VFs, i.e., the case for Δ*x*_*L*_ = Δ*x*_*R*_ = 0. The collision height is calculated simply based on the collision condition and the length of the VF:

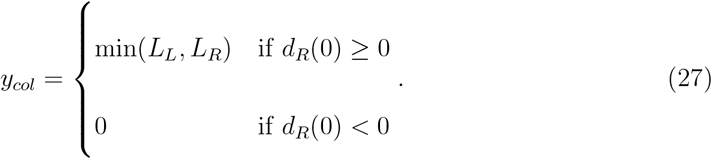

The effective spring collision constant in Equation 19 is computed assuming an in-series spring configuration:

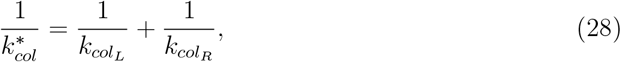

which comprises the contributions from the left- and right-side collision springs. The values for 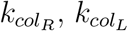 and *η*_*col*_ are computed following Galindo *et al*. (2017).

Replacing the Eqs. (23) and (28) in the integral in Eq. (19) the total collision force can be computed by:

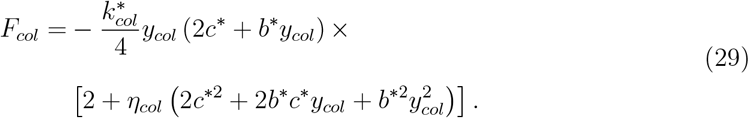

Note that the quantities 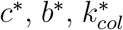, and *y*_*col*_ have information on the properties of both VFs.

## Notes

### Competing Interest Statement

The authors have declared no competing interest.

## References

Alzamendi, G. A., Peterson, S. D., Erath, B. D., Hillman, R. E., and Zañartu, M. (2022). “Triangular body-cover model of the vocal folds with coordinated activation of the five intrinsic laryngeal muscles,” The Journal of the Acoustical Society of America 151(1), 17–30.

Birkholz, P. (2016). “Glottalimageexplorer an open source tool for glottis segmentation in endoscopic high-speed videos of the vocal folds,” in Studientexte zur Sprachkommunikation: Elektronische Sprachsignalverarbeitung 2016, edited by O. Jokisch (TUDpress, Dresden), pp. 39–44.

Birkholz, P., Kröger, B. J., and Neuschaefer-Rube, C. (2011). “Synthesis of breathy, normal, and pressed phonation using a two-mass model with a triangular glottis,” in Interspeech 2011: 12th Annual Conference of the International Speech Communication Association, pp. 2681–2684, Florence, Italy.

Bonilha, H. S., Deliyski, D. D., and Gerlach, T. T. (2008). “Phase asymmetries in normophonic speakers: Visual judgments and objective findings,” American Journal of Speech-Language Pathology 17(4), 367–376, 10.1044/1058-0360(2008/07-0059), doi:10.1044/1058-0360(2008/07-0059).

Chhetri, D. K., Berke, G. S., Lotfizadeh, A., and Goodyer, E. (2009). “Control of vocal fold cover stiffness by laryngeal muscles: A preliminary study,” Laryngoscope 119(1), 222–227.

Desjardins, M., Apfelbach, C., Rubino, M., and Abbott, K. V. (2022). “Integrative review and framework of suggested mechanisms in primary muscle tension dysphonia,” Journal of Speech, Language, and Hearing Research 65(5), 1867–1893, https://doi.org/10.1044/5072022_jslhr-21-00575,, doi:10.1044/2022_jslhr-21-00575.

Dresel, C., Mergell, P., Hoppe, U., and Eysholdt, U. (2006). “An asymmetric smooth contour two-mass model for recurrent laryngeal nerve paralysis,” Logopedics Phoniatrics Vocology 31(2), 61–75, https://doi.org/10.1080/14015430500363232, doi:10.1080/14015430500363232.

Erath, B., Zañartu, M.,, Peterson, S., and Plesniak, M. (2011). “Nonlinear vocal fold dynamics resulting from asymmetric fluid loading on a two-mass model of speech,” Chaos 21, 033113–1, doi:10.1063/1.3615726.

Erath, B., Zañartu, M., Stewart, K., Plesniak, M., Sommer, D., and Peterson, S. (2013). “A review of lumped-element models of voiced speech,” Speech Communication 55, 667690, doi:10.1016/j.specom.2013.02.002.

Erath, B. D., Peterson, S. D., Weiland, K. S., Plesniak, M. W., and Zañartu, M. (2019). “An acoustic source model for asymmetric intraglottal flow with application to reduced-order models of the vocal folds,” PLoS ONE 14(7), 1–27.

Galindo, G. E., Peterson, S. D., Erath, B. D., Castro, C., Hillman, R. E., and Zañartu, M. (2017). “Modeling the pathophysiology of phonotraumatic vocal hyperfunction with a triangular glottal model of the vocal folds,” Journal of Speech, Language, and Hearing Research 60(9), 2452–2471.

Horáček, J., Laukkanen, A. M., and Šidlof, P. (2007). “Estimation of impact stress using an aeroelastic model of voice production,” Logopedics Phoniatrics Vocology 32(4), 185–192.

Hsiung, M.-W., and Hsiao, Y.-C. (2004). “The Characteristic Features of Muscle Tension Dysphonia before and after Surgery in Benign Lesions of the Vocal Fold,” ORL 66(5), 246–254.

Hunter, E. J., Titze, I. R., and Alipour, F. (2004). “A three-dimensional model of vocal fold abduction/adduction,” The Journal of the Acoustical Society of America 115(4), 1747–1759, 10.1121/1.1652033, doi:10.1121/1.1652033.

Ivey, C. M. (2019). “Vocal Fold Paresis,” Otolaryngologic clinics of North America 52(4), 637–648.

Jiang, J. J., Zhang, Y., and McGilligan, C. (2006). “Chaos in voice, from modeling to measurement,” Journal of Voice 20(1), 2–17.

Lohscheller, J., Eysholdt, U., Toy, H., and Dollinger, M. (2008). “Phonovibrography: Mapping high-speed movies of vocal fold vibrations into 2-d diagrams for visualizing and analyzing the underlying laryngeal dynamics,” IEEE Transactions on Medical Imaging 27(3), 300–309, 10.1109/tmi.2007.903690, doi:10.1109/tmi.2007.903690.

Lucero, J. C., and Koenig, L. L. (2005). “Simulations of temporal patterns of oral airflow in men and women using a two-mass model of the vocal folds under dynamic control,” The Journal of the Acoustical Society of America 117(3), 1362–1372.

Lucero, J. C., Pelorson, X., and Hirtum, A. V. (2019). “Vocal fold oscillators at large asymmetries,” Models and Analysis of Vocal Emissions for Biomedical Applications - 11th International Workshop, MAVEBA 2019 (December), 145–148.

Lucero, J. C., Pelorson, X., and Van Hirtum, A. (2020). “Phonation threshold pressure at large asymmetries of the vocal folds,” Biomedical Signal Processing and Control 62(July).

Lucero, J. C., and Schoentgen, J. (2015). “Smoothness of an equation for the glottal flow rate versus the glottal area,” The Journal of the Acoustical Society of America 137(5), 2970–2973, http://asa.scitation.org/doi/10.1121/1.4919297http://dx.doi.org/10.1121/1.4919297, doi:10.1121/1.4919297.

Manríquez, R., Peterson, S. D., Prado, P., Orio, P., Galindo, G. E., and Zanartu, M. (2019). “Neurophysiological Muscle Activation Scheme for Controlling Vocal Fold Models,” IEEE Transactions on Neural Systems and Rehabilitation Engineering 27(5), 1043–1052.

Mattheus, W., and Brucker, C. (2011). “Asymmetric glottal jet deflection: differences of two-and three-dimensional models.,” The Journal of the Acoustical Society of America 130(6), EL373–9.

Mehta, D. D., Deliyski, D. D., Quatieri, T. F., and Hillman, R. E. (2011). “Automated measurement of vocal fold vibratory asymmetry from high-speed videoendoscopy recordings,” Journal of speech, language, and hearing research : JSLHR 54(1), 47–54.

Pickup, B. A., and Thomson, S. L. (2009). “Influence of asymmetric stiffness on the structural and aerodynamic response of synthetic vocal fold models,” Journal of Biomechanics 42(14), 2219–2225.

Qiu, Q., Schutte, H., Gu, L., and Yu, Q. (2003). “An automatic method to quantify the vibration properties of human vocal folds via videokymography,” Folia Phoniatrica et Logopaedica 55(3), 128–136, 10.1159/000070724, doi:10.1159/000070724.

Samlan, R. A., and Story, B. H. (2017). “Influence of LeftRight Asymmetries on Voice Quality in Simulated Paramedian Vocal Fold Paralysis,” Journal of Speech, Language, and Hearing Research 60(2), 306–321.

Samlan, R. A., Story, B. H., Lotto, A. J., and Bunton, K. (2014). “Acoustic and perceptual effects of left–right laryngeal asymmetries based on computational modeling,” Journal of Speech, Language, and Hearing Research 57(5), 1619–1637, https://doi.org/10.1044/5762014_jslhr-s-12-0405, doi:10.1044/2014_jslhr-s-12-0405.

Smith, B. L., Nemcek, S. P., Swinarski, K. A., and Jiang, J. J. (2013). “Nonlinear source-filter coupling due to the addition of a simplified vocal tract model for excised larynx experiments,” Journal of Voice 27(3), 261–266.

Sommer, D. E., Erath, B. D., Zañartu, M., and Peterson, S. D. (2013). “The impact of glottal area discontinuities on block-type vocal fold models with asymmetric tissue properties,” The Journal of the Acoustical Society of America 133(3), EL214–EL220, https://doi.org/10.1121/1.4790662, doi:10.1121/1.4790662.

Spencer, M. L. (2015). “Muscle Tension Dysphonia: A Rationale for Symptomatic Subtypes, Expedited Treatment, and Increased Therapy Compliance,” Perspectives on Voice and Voice Disorders 25(1), 5–15.

Steinecke, I., and Herzel, H. (1995). “Bifurcations in an asymmetric vocal-fold model,” Journal of the Acoustical Society of America 97(3), 1874–1884.

Story, B. H. (2002). “An overview of the physiology, physics and modeling of the sound source for vowels,” Acoustical Science and Technology 23(4), 195–206.

Story, B. H. (2008). “Comparison of magnetic resonance imaging-based vocal tract area functions obtained from the same speaker in 1994 and 2002,” The Journal of the Acoustical Society of America 123(1), 327–335.

Story, B. H. (2015). “Mechanisms of Voice Production,” The Handbook of Speech Production (January), 34–58.

Story, B. H., and Titze, I. R. (1995). “Voice simulation with a body-cover model of the vocal folds,” Journal of the Acoustical Society of America 97(2), 1249–1260.

Tao, C., and Jiang, J. J. (2007). “Mechanical stress during phonation in a self-oscillating finite-element vocal fold model.,” Journal of biomechanics 40(10), 2191–2198.

Tipton, P. W., Ekbom, D. C., Rutt, A. L., and van Gerpen, J. A. (2020). “Vocal Fold “Paralysis”: An Early Sign in Multiple System Atrophy,” Journal of voice 34(6), 940–944.

Titze, I. R. (2006). The Myoelastic Aerodynamic Theory of Phonation, 1st edition ed. (National Center for Voice and Speech).

Titze, I. R., and Hunter, E. J. (2007). “A two-dimensional biomechanical model of vocal fold posturing,” The Journal of the Acoustical Society of America 121(4), 2254.

Titze, I. R., and Story, B. H. (2002). “Rules for controlling low-dimensional vocal fold models with muscle activation,” The Journal of the Acoustical Society of America 112(3), 1064–1076.

Vampola, T., Horáček, J., and Klepáček, I. (2016). “Computer simulation of mucosal waves on vibrating human vocal folds,” Biocybernetics and Biomedical Engineering 36(3), 451–465.

Xue, Q., Mittal, R., Zheng, X., and Bielamowicz, S. (2010). “A computational study of the effect of vocal-fold asymmetry on phonation,” The Journal of the Acoustical Society of America 128(2), 818–827, 10.1121/1.3458839, doi:10.1121/1.3458839.

Yokota, K., Ishikawa, S., Koba, Y., Kijimoto, S., and Sugiki, S. (2019). “Inverse analysis of vocal sound source using an analytical model of the vocal tract,” Applied Acoustics 150, 89–103.

Zañartu, M. (2006). “Influence of acoustic loading on the flow-induced oscillations of single mass models of the human larynx,” Master’s thesis, School of Electrical and Computer Engineering, Purdue University, West Lafayette, IN.

Zañartu, M. (2010). “Acoustic coupling in phonation and its effect on inverse filtering of oral airflow and neck surface acceleration,” Ph.D. thesis, School of Electrical and Computer Engineering, Purdue University, West Lafayette, IN.

Zañartu, M., Galindo, G. E., Erath, B. D., Peterson, S. D., Wodicka, G. R., and Hillman, R. E. (2014). “Modeling the effects of a posterior glottal opening on vocal fold dynamics with implications for vocal hyperfunction,” The Journal of the Acoustical Society of America 136(6), 3262–3271.

Zhang, L., Li, J., Dong, M., Cui, Y., and Rong, X. (2019). “Development of a Compatible Exoskeleton (Co-Exos II) for Upper-Limb Rehabilitation,” Institute of Electrical and Electronics Engineers Inc., pp. 240–245.

Zhang, Y., and Jiang, J. J. (2004). “Chaotic vibrations of a vocal fold model with a unilateral polyp,” The Journal of the Acoustical Society of America 115(3), 1266–1269.

Zhang, Z. (2016). “Cause-effect relationship between vocal fold physiology and voice production in a three-dimensional phonation model,” The Journal of the Acoustical Society of America 139(4), 1493–1507.

Zhang, Z. (2018). “Vocal instabilities in a three-dimensional body-cover phonation model,” The Journal of the Acoustical Society of America 144(3), 1216–1230.

Zhang, Z., and Luu, T. H. (2012). “Asymmetric vibration in a two-layer vocal fold model with left-right stiffness asymmetry: Experiment and simulation,” The Journal of the Acoustical Society of America 132(3), 1626–1635, 10.1121/1.4739437, doi:10.1121/1.4739437.

